# A microtubule-based mechanism predicts cell division orientation in plant embryogenesis

**DOI:** 10.1101/270793

**Authors:** Bandan Chakrabortty, Viola Willemsen, Thijs de Zeeuw, Che-Yang Liao, Dolf Weijers, Bela Mulder, Ben Scheres

## Abstract

Oriented cell divisions are significant in plant morphogenesis because plant cells are embedded in cell walls and cannot relocate. Cell divisions follow various regular orientations, but the underlying mechanisms have not been clarified. We show that cell-shape dependent self-organisation of cortical microtubule arrays is crucial for determining planes of early tissue-generating divisions and forms the basis for robust control of cell division orientation in the embryo. To achieve this, we simulate microtubules on actual cell surface shapes from which we derive a minimal set of three rules for proper array orientation. The first rule captures the effects of cell shape alone on microtubule organisation, the second rule describes the regulation of microtubule stability at cell edges and the third rule includes the differential effect of auxin on local microtubule stability. These rules explain early embryonic division plane orientations and offer a framework for understanding patterned cell divisions in plant morphogenesis.

## Introduction

The striking regularities in plant cell division patterns have spurred the formulation of heuristic geometric rules for oriented cell divisions for already more than a century. These rules relate division planes e.g. to the principal direction of growth, geometric relations to existing cell walls and the nucleus, or minimum cell surface energy[1–6]. With the advent of modern cell biology, division plane orientation in plants has been correlated to the orientation of the ensemble of microtubules (MTs) underneath the plasma membrane, the cortical microtubule array (CMA)[7,8]. Shortly before cell division, the MTs get restricted to a plane that is closely associated with the nucleus, forming the so-called pre-prophase band (PPB). PPB and, by extension, CMA orientation is an indicator of cell division orientation[9,10]. Some variability between PPB and cell plate orientation may occur (e.g. Oud and Nanninga[11]) and recent evidence suggests that the PPB is not strictly required for division plane orientation[12], but nevertheless the link of cell division plane to CMA orientation is maintained. As cortical MTs co-align with cellulose microfibrils, growing cells exhibit a complex interplay between CMA orientation, the deposition of microfibrils in the cell wall and growth[13,14]. There is evidence that MT severing proteins may differentially act on MTs under tension, and cell growth and cell wall anisotropy may, through this mechanism, orient the CMA perpendicular to the cell growth axis[15] ‒underpinning the oldest proposed geometric rule^1^. However, many formative divisions, which create ordered tissue layers, take place in regions of limited growth. A prime example is the early stage embryo in *Arabidopsis*[16–18]. Previous analysis of *Arabidopsis* embryogenesis shows that auxin-insensitive *bodenlos* (*bdl*) embryos divide according to the proposed geometric rules, whereas wild-type (WT) divisions seem to require additional control[16], possibly through auxin-mediated CMA regulation. Neither the molecular mechanisms underlying the phenomenological geometric rules nor the nature of the required additional control is understood, hence the field lacks models to explain the orientation of formative cell divisions. To establish such models, factors must be identified that influence the ordering of the CMA in volumetrically slowly growing cells and are based on the fundamental processes driving cortical MT ensembles, such as the dynamical instability mechanism[19] and MT-MT collisions[20]. Here, we identify candidate molecular processes for cell division orientation control in embryos by utilizing a comprehensive computational framework that simulates MT ordering on slow-growing cell surfaces with realistic cell shapes. Combining the in-silico results with high-resolution visualization of both cell-outline and cortical MTs in WT *Arabidopsis* embryos and selected mutants enables us to show that hitherto unanticipated constraints due to cell shape, in combination with cell edge-catastrophe protection and auxin-mediated MT stability suffice to explain cell division patterns in 1- to 16-cell stage embryos. Our MT-based molecular statistical mechanism predicts division plane orientations, recapitulates many of the proposed geometric rules and establishes a basis for specific experimental validations in the future.

## Results

### First principle-based MT modelling on realistic cell shapes

Our approach for understanding rules of cell division in cells that undergo formative cell divisions was to model cortical MT dynamics during the first four division cycles in the *Arabidopsis* embryo proper, which have been documented in detail[16,21–23]. These characteristic divisions lead to the formation of inner and outer tissue layers (Figure 1(A), left). Our starting assumption was that embryonic cells, like most cell types previously considered, have a regular CMA orientation that predicts the orientation of cell division. To test this assumption, we generated transgenic lines and optimized imaging methodology for high-resolution 3D-imaging of the CMA throughout the different stages of *Arabidopsis* embryogenesis. We expressed TUA6-GFP from the embryo specific *WUSCHEL RELATED HOMEOBOX 2* (*WOX2*) promoter. Maximum projections of high-resolution Z-stacks reveal the overall topology of the CMA for embryos from the 1-cell stage to globular stage (Figure 1(A), right). Cell segmentation allowed the extraction of individual cell surfaces for the different embryonic cells. Analysis of cortical MT signals projected on these extracted cell surfaces shows the ordered orientation of CMA on individual cells that collapse to form PPB structures (Figure 1(B)) which correctly predict future division plane orientation (Figure 1(C)). Our observations validated the early embryo as a suitable model system to investigate how cortical MT ordering is established in cells with varying geometries.

**Figure 1.**
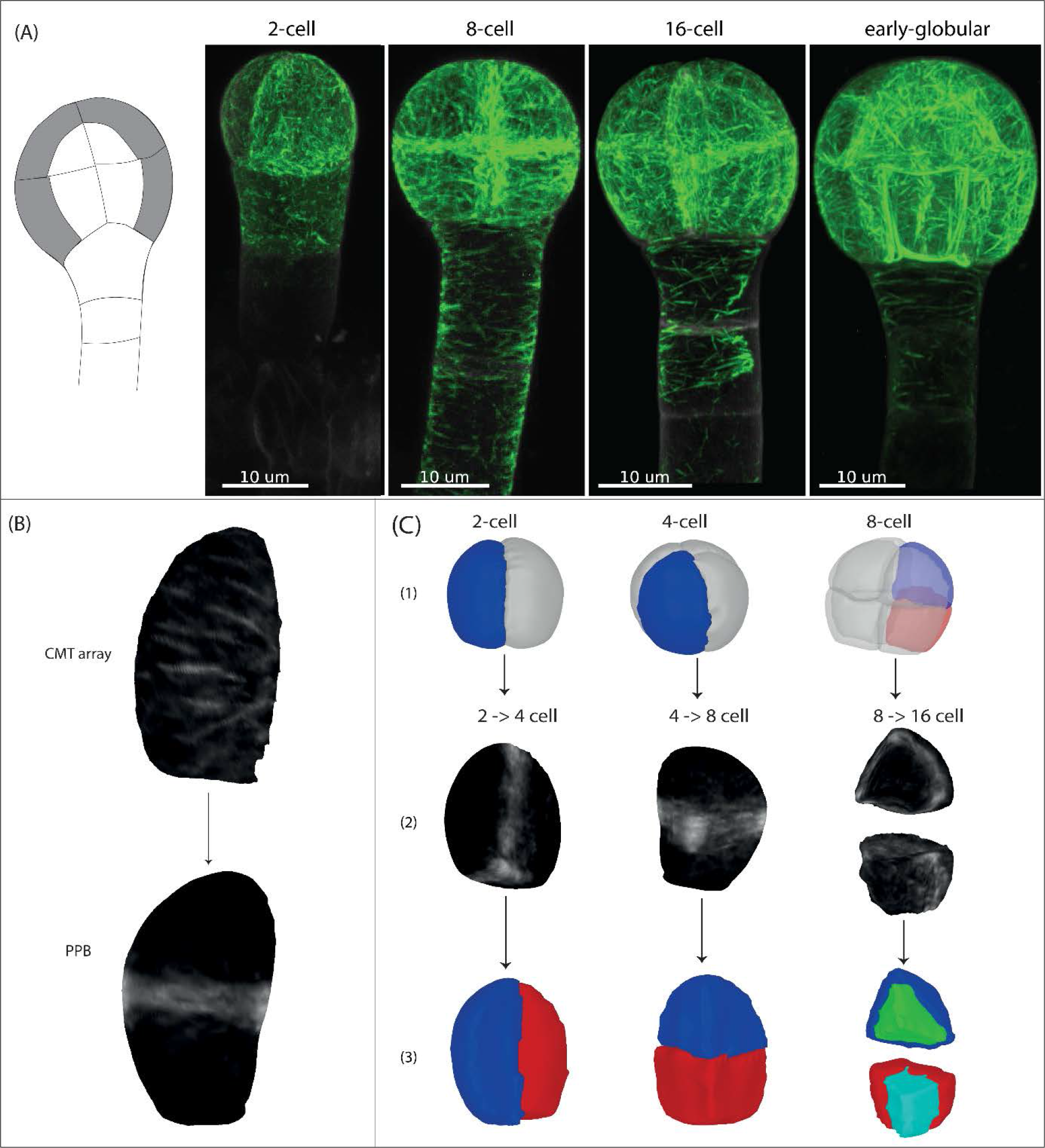
Experimental visualization of CMA and PPB during *Arabidopsis* early embryonic development. (A) The characteristic divisions lead to the formation of inner and outer tissue layers (left), cortical MTs visualized using a pWOX2::TUA6-GFP reporter line show an ordered orientation of CMA (right). (B) Sequential imaging of cortical MTs and the SR2200 membrane stain allows for extraction of cell shapes for separate embryonic cells. Projection of cortical MT signal on extracted cell shapes shows orientation of CMA that predicts the orientation of PPB (7 cells were analysed). (C) PPB predicts the location and orientation of the division plane: (1) Representative single cell from 2-cell stage and 4-cell stage embryo, and representative single cell from upper tier and lower tier of 8-cell stage embryo are highlighted, (2) Projection of PPB on the corresponding cell templates during 2- to 4-cell stage (3 cells were analysed), 4- to 8-cell stage (5 cells were analysed) and 8- to 16-cell stage (5 cells were analysed) transitions, and (3) Pair of daughter cells during 2- to 4-, 4- to 8- and 8- to 16-cell stage transitions.

To enable the systematic exploration of factors that govern CMA orientation in the early embryo, we designed a computational framework to simulate the key properties of MT dynamics on arbitrary shaped cellular surfaces. CMAs in animal cells have been shown to be sensitive to cell shape[24]. Therefore, we used the actual embryonic cell shapes obtained by segmenting high-resolution confocal images of fluorescently stained *Arabidopsis* embryos to extract cell surfaces, approximated through fine-grained triangulation (Figure 2 (A)). To simulate MT dynamics on individual planar triangles we adapted a previously developed event-driven algorithm[25,26] that implements all key MT properties with biophysical significance, such as nucleation, growth, catastrophe and rescue rates. Three possible results from MT-MT collisions—zippering, crossover and induced-catastrophe—are also encoded (Figure 2(B)). Although additional factors are known to influence MT dynamics, most notably directional nucleation from existing MTs[27] and severing by katanin [28,29], previous modelling work has shown that these additional effects mainly modify the range of parameters for which ordered arrays develop spontaneously, but have less impact on the nature of the ordered state per se [30–32]. Parsimony therefore led us to omit these features here. The trajectories of the simulated MT segments on the individual triangles are ‘glued’ together consistent with the 3-dimensional cell geometry employing the connectivity graph of the triangulated cell surface[33] (Figure 2(D)). All parameters of individual MT behaviour, excluding collision parameters, can be absorbed in a single control parameter (G) that controls the frequency of MT interactions[31,34]. If G reaches a threshold value interacting cortical MTs will spontaneously form an ordered array. The interpretation of changing G is flexible, as equivalent changes in G can be brought about by different changes in the underlying biophysical parameters. Our ability to tag triangles belonging to particular cell faces and cell edges enabled us to define regions on which we could independently vary the G parameter. Finally, to read out the order of the steady-state CMA, we employed a tensorial order parameter **Q**^(2)^, whose absolutely smallest eigenvalue Q^(2)^ characterizes the degree of ordering: Q^(2)^ = 0 when MT orientation is random and Q^(2)^ ~ 1 when MTs are completely ordered. The normalized eigenvector 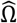 associated with Q^(2)^ characterizes the global orientation of the MT array and is, in all simulation images, represented by a dot marking the end point of the vector[33] (Figure 2(C)).

**Figure 2.**
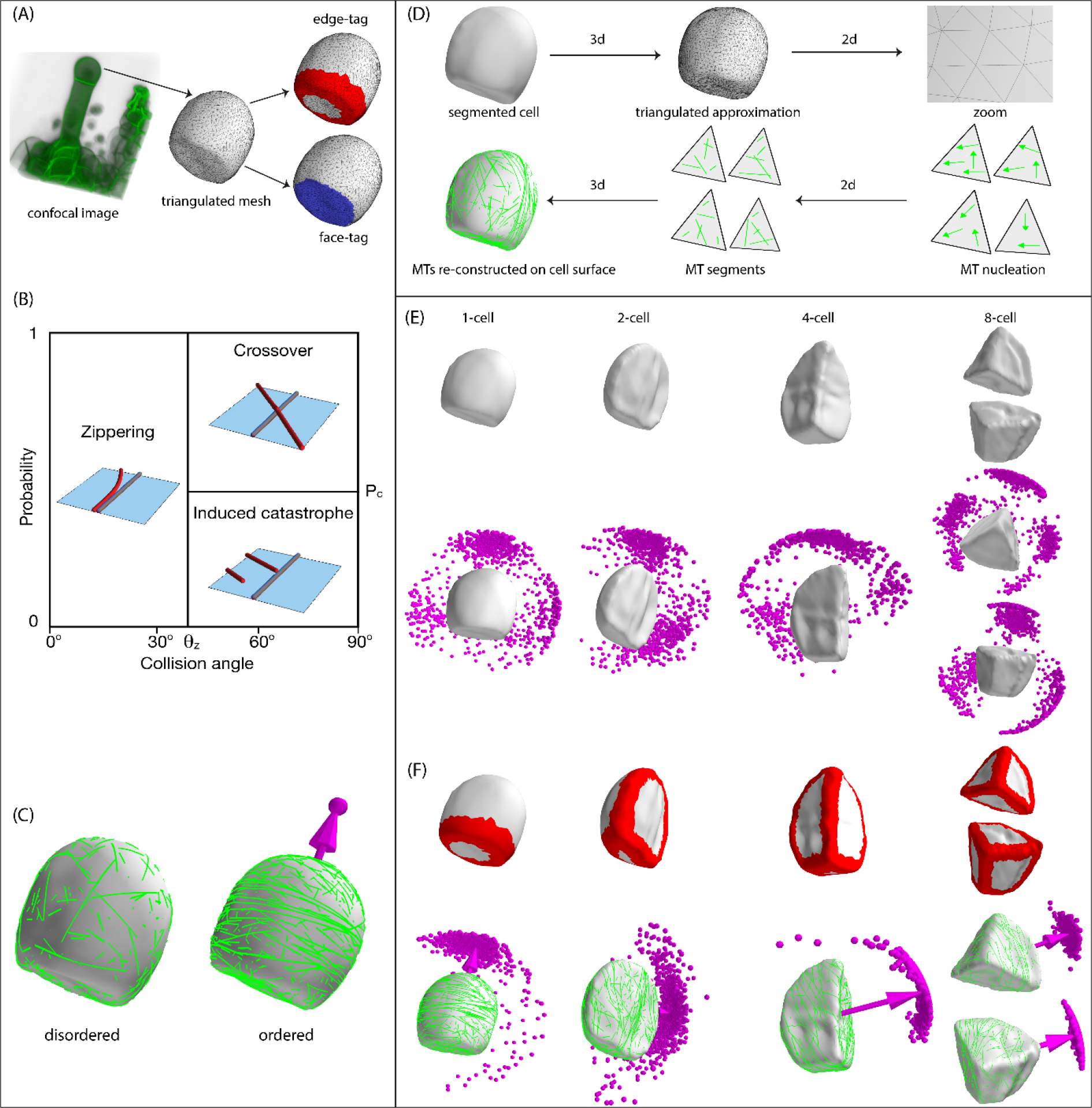
MT modelling on *Arabidopsis* early embryonic cell templates. (A) The confocal image of embryonic 1-cell shape is extracted as a triangulated surface mesh via image segmentation software MorphoGraphX[46]. The edge of the mesh is tagged (red colour) by assigning all triangles belonging to this edge with a unique edge-tag index. The flat bottom face (faint blue) and the curved top face (faint grey) are tagged by assigning all their triangles with a unique face-tag index. Edge-tags and face-tags are used for simulation implementation of edge-catastrophe and face specific MT stabilization. (B) Basic MT-MT interactions. Two dimensional attachment of MTs to cell cortex allows MTs to interact with each other via collisions. Shallow angles (≤ 40°) of collision lead to zippering and steeper angles (≥ 40°) of collision lead to crossover or induced-catastrophe. (C) Order parameter to quantify the degree of MT order and array orientation. A disordered state results in Q^(2)^ = 0 without formation of MT array. An ordered state indicates the formation of MT array with significant degree of MT order, Q^(2)^ ~ 1. Perpendicular to the MT array, a vector 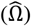 is defined which quantifies the orientation of the MT array. The tip of this vector will be used to represent the MT array orientation. (D) A schematic overview of the simulation approach, where geometrically correct propagation of growing MT ends between the neighbouring triangles lead to the reconstruction of MT dynamics on the cell surfaces. Simulated orientation of MT array on 1- to 8-cell stage WT cell templates: (E) Default cell shapes, which showed a diverse but not randomly distributed MT array orientation, and (F) With edge-catastrophe in MT dynamics, which showed one unique cluster of MT array orientation in each cell stage. Simulations were performed for ≈ 1000 independent configurations of stochastic MT dynamics. The arrow vectors represent mean orientation of the associated MT array.

### Shape and edge-catastrophe can explain early embryo cell division patterns in an auxin response mutant

We performed ≈ 1000 stochastically independent simulations of MT dynamics per cell on cell surfaces extracted from WT embryos from 1- to 8-cell stage, ensuring throughout that a steady state was reached. The resulting CMAs displayed a diverse but not randomly distributed orientation (Figure 2(E)). Thus, cell shape significantly impacted, but could not uniquely specify the orientation of the CMAs on a given cell template. It has been reported that cell edges may hinder the propagation of MTs in plant cells depending on their degree of curvature[35]. We were curious whether this reported property of MT behaviour could further reduce the diversity of CMA orientations in our simulations. We implemented a local enhanced catastrophe in those cell edges that have high curvature. In the various WT cells, this implementation robustly reduced the possible CMA orientations to a single sharply defined cluster for each cell stage (Figure 2(F)). However, the resultant inferred division patterns were non-WT and instead appeared to match those of the auxin insensitive *bdl* mutant[16] division patterns. Moreover, the simulations predicted a preference for slightly oblique division at the 4-cell stage in *bdl* mutant hitherto not described. We experimentally verified this preference of oblique division for 4- to 8-cell stage transition in the *bdl* mutant as well as their aberrant divisions in 8- to 16-cell stage transition[16] (Figure S1). Note that, at the 1- to 2-cell transition, all simulations yielded horizontal divisions, whereas only 18% of *bdl* mutants divide in this direction[36]. To exclude that the mapping to *bdl* division patterns was a coincidental effect of using WT cell templates, we extracted *bdl* cell templates from 4- to 16-cell stages, reconstructed progenitor 2- to 8-cell stage cell surfaces by merging corresponding daughter cell pairs, and ran simulations on these reconstructed ‘mother cells’ (Figure 3(A)). The steady state CMA orientation in all these simulations correctly predicted the division plane, confirming that shape effects and edge-catastrophe were sufficient to explain *bdl* divisions between 2- to 16-cell stages (Figure 3(B)).

**Figure 3.**
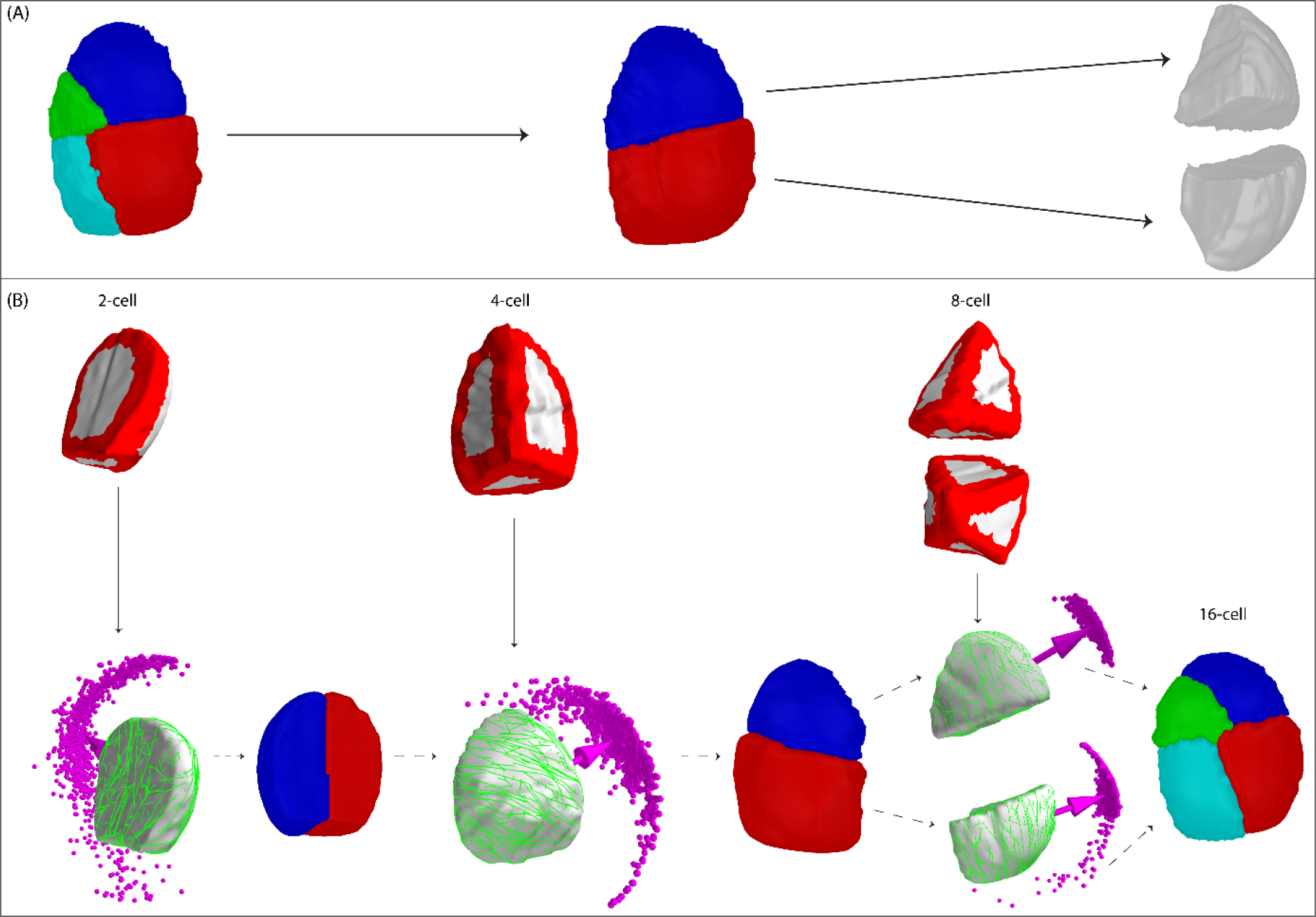
Recapitulation of *bdl* division patterns in *Arabidopsis* early embryonic development. (A) Reconstruction of mother cell by merging the corresponding daughter cell pair. In this example, the daughter pair of both upper and lower tier of *Arabidopsis bdl* 16-cell template were merged to reconstruct the corresponding mother 8-cell template. Such re-created mother cells were separated out for simulating MT dynamics. (B) Simulated MT arrays on 2- to 8-cell stage of *bdl* cell template, which were reconstructed by merging the corresponding daughter cell pair of the next cell stage. Simulations were performed with edge-catastrophe, yielding one unique cluster of MT array orientation in each cell stage, which correctly predicted the corresponding *bdl* division plane orientation. Cell edges are coloured red to indicate that in simulation MTs were subjected to edge-catastrophes.

### Face stability and edge-catastrophe reduction explain WT early embryo cell division patterns

The highly characteristic WT division pattern at the 8- to 16-cell stage transition that separates inner and outer cell layers did not follow the rules based on cell shape and edge-catastrophe only. To search for an additional rule that might explain the WT division pattern, we first focused on the divisions between 2- to 16-cell stages, which were correctly predicted in the *bdl* mutant. We asked whether the observed defects with respect to WT were associated with aberrant CMA formation. Upon close inspection of CMAs in *RPS5A>bdl*[16] embryos, ubiquitously expressing mutant bdl protein, MTs were shorter on cell faces compared to WT, indicating that reduced auxin signalling affects MT length in embryo cells (Figure S2). We implemented this observation in our simulations by allowing auxin-dependent cell face-specific changes in the G parameter, leading to locally enhanced average MT length (Figure S3). The resulting simulations therefore implemented, next to the basic rules for MT dynamics, effects of two biological control parameters: auxin-regulated face stability (Figure 4(A)) and edge-catastrophe. We investigated multiple combinations of the strength of these two effects yielding different behavioural regimes. A complete match with WT division patterns (Figure 4(B), Figure S4) was obtained by simulating MT dynamics on ‘original’ or ‘reconstructed mother’ 2- to 8-cell stage cell templates under the following adaptations of our initial rules: (1) enhanced cell face stability that is strongest in recent division faces and (2) reduced edge-catastrophe (for simulation parameters see Table S1). These specific rules conceivably result from concrete molecular processes. First, auxin-mediated enhanced face stability could be transiently established at each new division site. Second, edge-catastrophe could be reduced, for example, by the activity of proteins that stabilize bending MTs[35].

**Figure 4.**
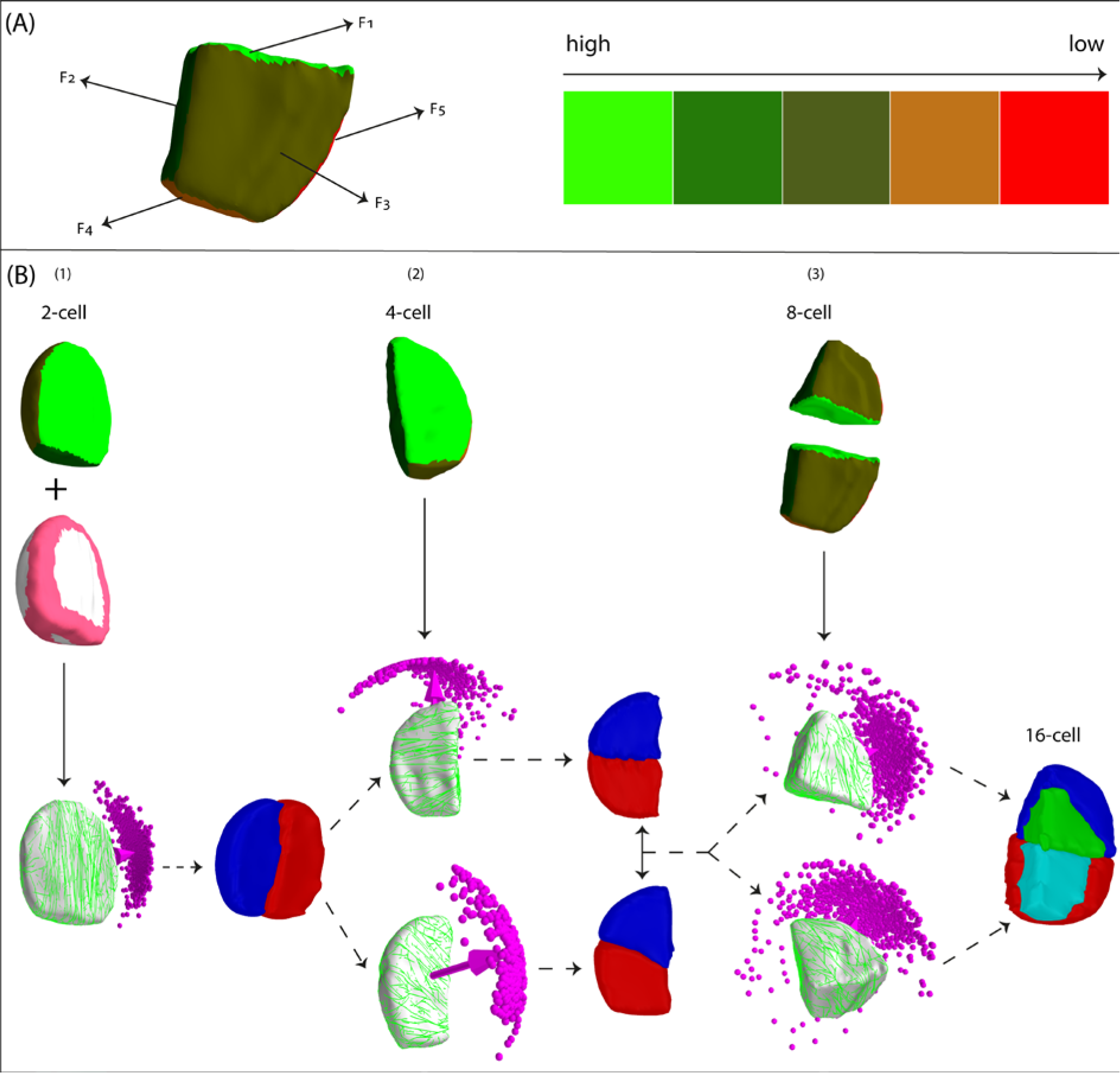
WT division pattern in *Arabidopsis* early embryonic development. (A) Developmental age of cell faces F_5_ > F_4_ > F_3_ > F_2_ > F_1_ as depicted by the colour scale. MT dynamic catastrophe rate increases with developmental age, yielding a corresponding decrease in average MT length i.e. lower MT stability (see also Figure S3). (B) Simulated orientation of MT array on WT cell templates taken from different cell stages: (1) 2-cell stage with reduced edge-catastrophe in MT dynamics and enhanced MT stabilization at developmentally new cell faces, resulting in one unique cluster of MT array orientation correctly predicting the WT division plane orientation, (2) 4-cell stage without edge-catastrophe in MT dynamics and enhanced MT stabilization at developmentally new cell faces, resulting in two clusters of MT array orientation correctly predicting the two observed phenotypes of WT division plane orientation, and (3) 8-cell stage without edge-catastrophe in MT dynamics and enhanced MT stabilization at developmentally new cell faces, resulting in one unique cluster of MT array orientation for both upper and lower tier cells correctly predicting the WT division plane orientation.

As an initial test of the requirement of edge-catastrophe reduction for obtaining correctly oriented CMAs between 2- to 16-cell stages in WT, we analysed embryos homozygous for two different mutations in the *CLASP* gene, reported to influence MT edge-catastrophe[35]. To support our assumption that edge-catastrophe is regulated in WT embryos by CLASP, its mutations should affect division plane orientation in a manner predicted by our simulations. Indeed, in both mutant alleles, division orientations were severely skewed compared to WT (Figure 5(A) for 4- to 8-cell stage; Figure S5 for all 1- to 16-cell stage). To test whether we could correctly predict division planes in *clasp* mutant embryos, we first focused on the resulting sister cells of 4- to 8-cell stage transition in the *clasp* mutant to reconstruct mother cell and then simulated CMA orientation on the resulting cell shape. We used WT settings for auxin-mediated face stability, assumed to be unaffected, but introduced increased edge-catastrophes to simulate loss of CLASP. Under these conditions, the observed CMA orientations indeed robustly predicted the observed division planes in the *clasp* mutant (Figure 5(B)). In conclusion, the model assumptions that result in WT division pattern also yield, mutatis mutandis, the aberrant division pattern of *clasp* at the critical 4- and 8-cell stage transition.

**Figure 5.**
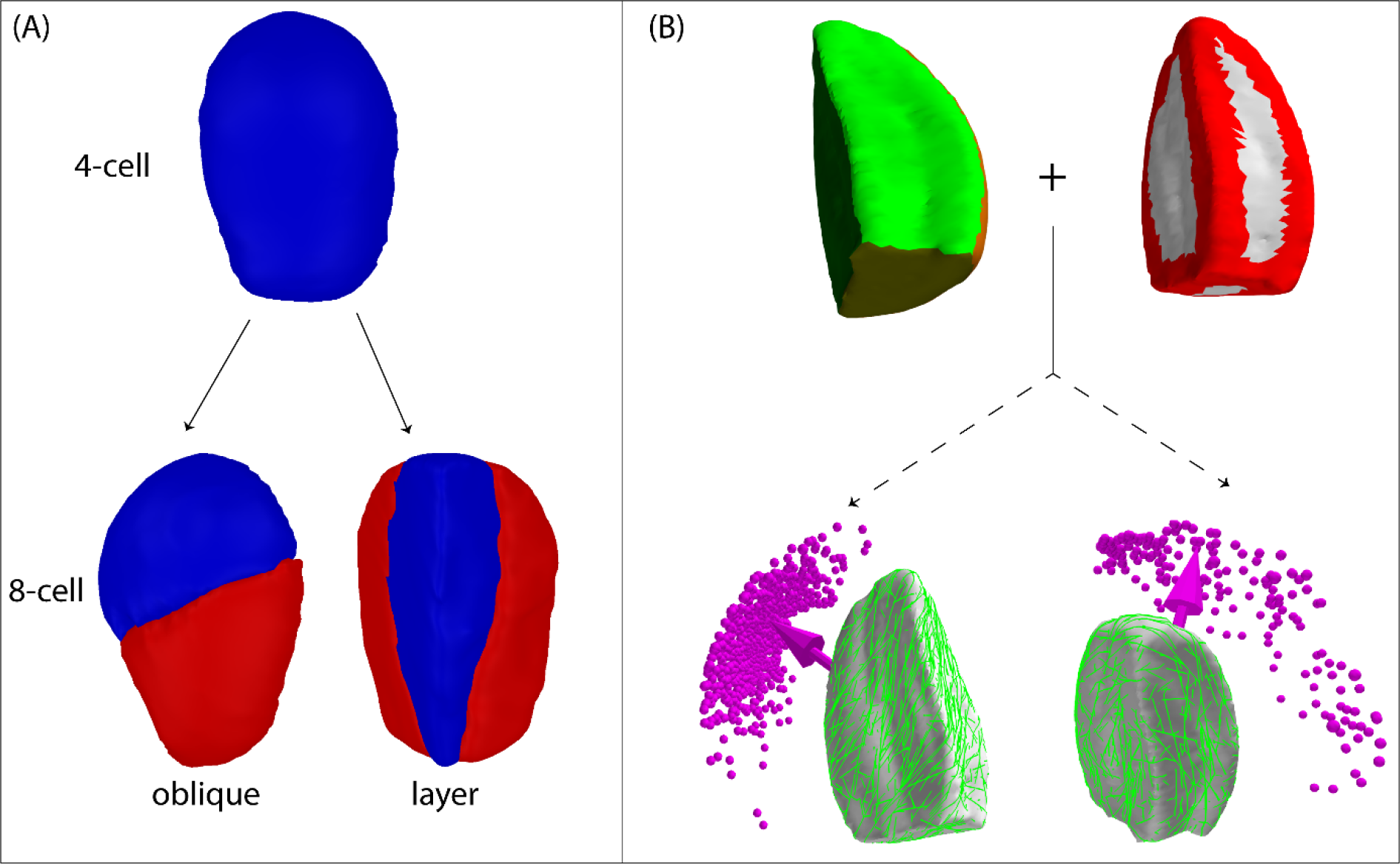
Simulating *clasp* mutant division patterns in *Arabidopsis* early embryonic development. (A) Representative *clasp* (*clasp 1* and *clasp 2*) division phenotypes during 4- to 8-cell stage transition (detailed quantification in Figure S5). (B) Simulated MT arrays on *clasp* 2 cell template of 4-cell stage, reconstructed by merging the corresponding 8-cell daughter cell pair. Simulations were performed with edge-catastrophe and enhanced MT stabilization at developmentally new cell faces, yielding two clusters of MT array orientation correctly predicting the two *clasp* division phenotypes. Cell edges coloured red indicate presence of edge-catastrophes and different colour coding of the cell faces represents the degree of MT stabilization (see Figure S3).

### Tuning of face stability and edge-catastrophe allows prediction of all early embryo cell division patterns

Our model so far predicted correct WT division patterns between the 2- to 16-cell stage when both auxin-mediated face stability and CLASP-mediated edge-catastrophe reduction are implemented. This same implementation of the model also correctly predicted the vertical orientation of the cell division plane in the 1- to 2-cell transition, albeit not fixing its in-plane direction (Figure S6(A) for WT cell template and Figure S6(B) for *clasp* cell template). Notably, even in the presence of edge-catastrophe (simulated as the *clasp* mutant) the correct vertical orientation is maintained (Figure 6(A) for WT cell template and Figure 6(B) for *clasp* cell template). Moreover, with moderate edge-catastrophe the division axis was constrained to a single specific direction, reminiscent of the true WT situation. This led us to hypothesise that CLASP is less active at the 1-cell stage than at later stages. Strikingly, when we implemented this stage-specific reduced CLASP activity without the auxin-mediated face stability rule (i.e., simulating a *bdl* mutant under this new condition), we obtained a proportion of horizontal and vertical divisions that closely matched the 18% horizontal divisions observed in *bdl* mutants (Figure 6(C)). In conclusion, auxin-mediated face stability from the 1-cell stage onward and CLASP-mediated edge-catastrophe reduction from the 2-cell stage onward can robustly and consistently recapitulate cell division orientations between the 1- to 16- cell stage in WT and *bdl* embryos, and between 1- to 8-cell stage in *clasp* embryos.

**Figure 6.**
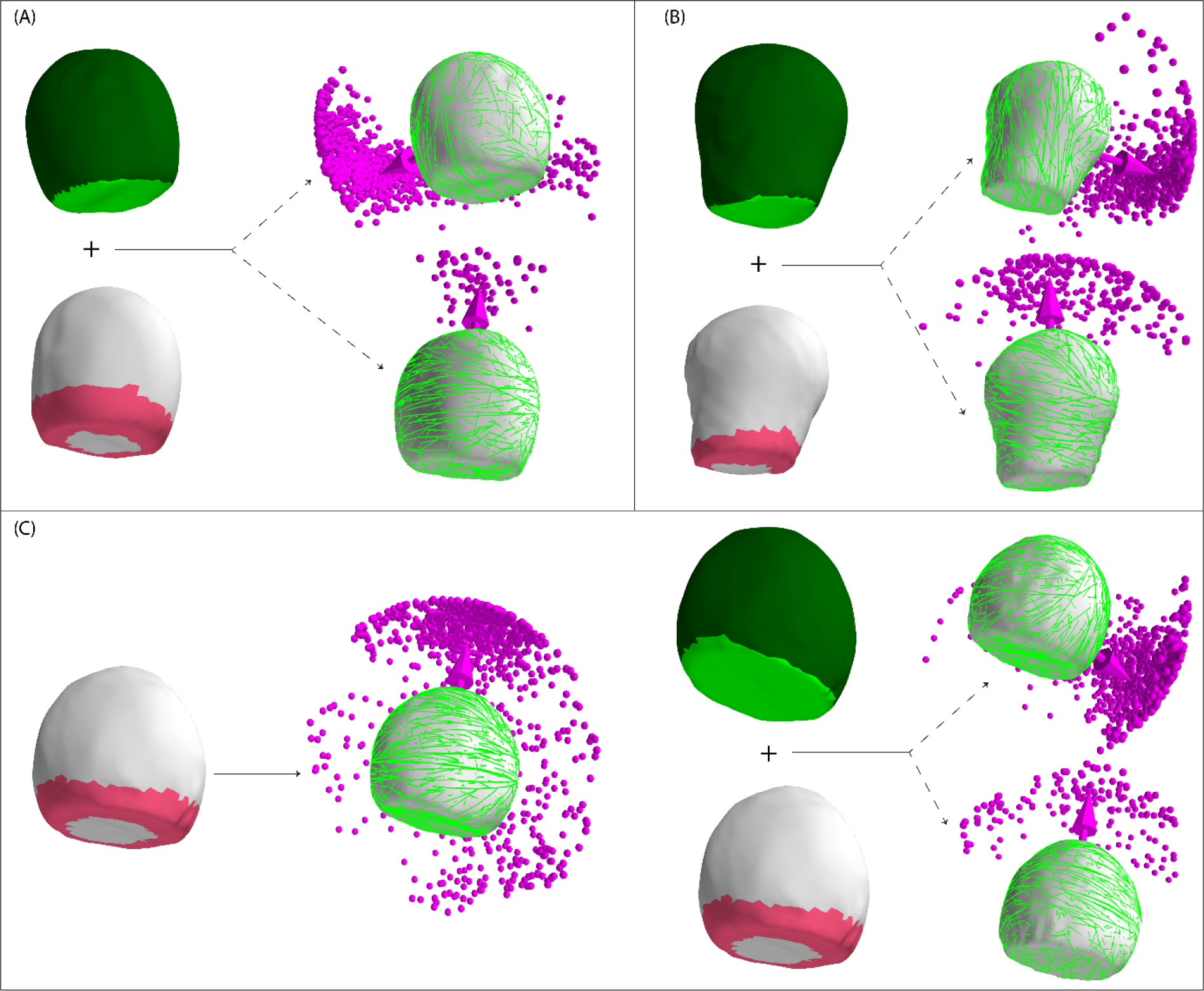
Fine-tuning of the 1- to 2-cell stage WT, *clasp* and *bdl* division patterns in *Arabidopsis* early embryos. (A) Simulations on WT cell template with reduced edge-catastrophe in MT dynamics and enhanced MT stabilization at developmentally new cell faces, showing a large cluster of a unique vertical MT array orientation (≈ 92%) that matched with the WT division plane orientation (top). A sparse cluster of horizontal MT array orientation (≈ 8%) was also observed (bottom). (B) Simulations on the clasp (*clasp* 2) cell template combining reduced edge-catastrophe and enhanced MT stabilization at developmentally new cell faces showed a majority cluster of vertical MT array orientations (≈ 75%) matching the *clasp* division plane orientation (top), and a sparse cluster of horizontal MT array orientations (≈ 25%), indicating stochasticity in division plane orientation under genetic perturbation (bottom). (C) Simulations on the *bdl* cell template with reduced edge-catastrophe resulted in a dominant horizontal orientation of MT array recapitulating the observed fraction (≈ 18%) of horizontal division plane orientation (left). Combination of reduced edge-catastrophe and enhanced MT stabilization at developmentally new cell faces produced a large cluster of MT array orientation (≈ 80%) which reflects the experimentally observed major vertical and unique division plane orientation (right, top) observed in *bdl* mutant. The remaining horizontal arrays (≈ 20%) indicate possible stochasticity in division plane orientation (right, bottom) under genetic perturbation, as actually observed and quantified^26^. Pink edges indicate reduced edge-catastrophes and colour coding of the cell faces represents the degree of MT stabilization (see Figure S3).

It is important to note that our quantification of the observed variation in the simulated CMA orientation (Figure 7) confirms the experimentally observed variation in division plane orientation in other cell types where formation PPB had been disturbed using *trm*678 mutant, i.e. epidermis, cortex and endodermis[12]. Finally, we noted that under these rules only a fraction of division predictions for the 8- to 16-cell transition of *clasp* mutants was correct. To assess this discrepancy, we looked at the growth rates of cells at this stage, inferred from the calculated cell volume increases between cell stages (Figure S6(D) and Figure S6(E)). We observed that the correctness of prediction negatively correlated with the growth rates of cell volume. Indeed, when, instead of using a re-created mother cell of the 4-cell stage embryo (by merging the corresponding daughter cell pair from 8-cell stage), we simulated MT dynamics on a true single cell of the 4-cell stage, the prediction of cell division plane orientation correlated better with the experimentally observed ones (Figure S6(C)). This result indicated that, in *clasp* mutants, cell growth becomes relevant. In this case, mechanisms that influence MT stability depending on direction of growth induced mechanical stress may become dominant factors in CMA orientation[37]. Hence, tissue growth rates define an upper bound below which the rules here described can successfully explain formative cell divisions.

**Figure 7.**
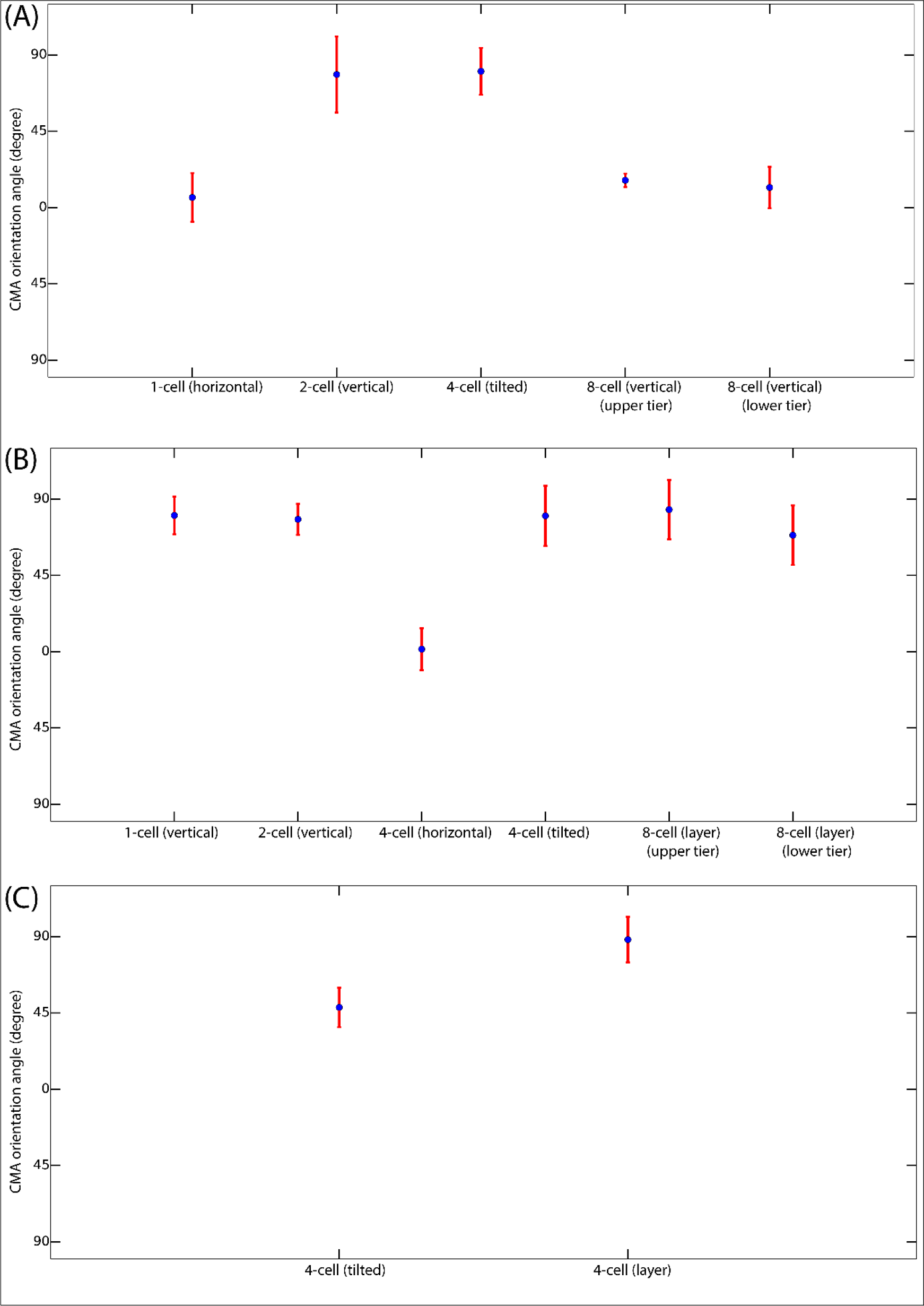
Quantification of the variation in simulated CMA orientation for individual cell stages of *Arabidopsis* early embryonic development: (A) *bdl* phenotype between 1- to 16-cell stage transitions, (B) WT phenotype between 1- to 16-cell stage transitions and (C) *clasp* phenotype between the critical 4- to 8-cell stage transition. CMA orientation angle was measured as the angle between the CMA array orientation vector and the longest principle axis of the cell for each cell stages. Simulated division phenotype is indicated in parentheses attached to different cell stage labelling.

## Discussion

Here we used a modelling framework on actual cell surface shapes and combined this with high-resolution imaging of both cell-outline and cortical MTs to derive a plausible model that explains the orientation of formative cell divisions in the *Arabidopsis* embryo from first principles on a molecular basis. When comparing predictions of division plane orientation based on basic MT-MT interaction rules and edge catastrophe with the predictions of the heuristic geometrical rules, it is striking that cell shape dependent MT based rules are by themselves already sufficient to correctly predict divisions in the auxin insensitive *bdl* mutant (Figure S7). The difference between *bdl* and WT division patterns indicates that, in addition, auxin-mediated regulation is crucial to establish the WT division pattern in *Arabidopsis* embryos. In order to explain the orientation of cell division up to the 16-cell stage of WT, we parsimoniously implemented, besides basic MT dynamics, two other factors influencing MT stability that are under biological control. The first biological control mechanism is an auxin-dependent face stability rule, derived from our observed MT patterns on *bdl* mutant embryos. The assumption that transient cortical MT stability is most strongly associated with new cell walls is not without precedence. Several observations indicate that peripheral marks instrumental for cell division orientation can transiently accumulate at specific locations in different organisms. In plants, *Arabidopsis* BASL marks are positioned away from the previous division wall[38]. In yeast, Bud8p and Bud9p proteins mark opposite division sites during bipolar budding[39]. The second biological control mechanism is a CLASP dependent edge-catastrophe reduction rule, based on published analyses of MTs at cell edges[35] and our own analysis of division planes in *clasp* mutant embryos. CLASP-like proteins in mammalian cells provide resistance to MTs under traction[40] and it is therefore conceivable that plant CLASPs in similar ways stabilize MT under torsion. It will be interesting to investigate whether mutations in transcription factors that specifically change early embryo division patterns, such as BDL[16] and WOX2[41], can be explained by their transcriptional control of MT regulators.

Our work provides a theoretical framework for fundamental understanding of formative cell divisions in plants, which are not restricted to embryogenesis but continuously occur in stem cell niches where tissue growth is reduced compared to surrounding zones (e.g.[17]). Live-cell imaging methods for Arabidopsis embryogenesis have recently been established[42,43]. Our model makes clear predictions on the key effects of auxin signalling and CLASP protein in specific cell faces and edges of the embryo and by extension other slow growing cells, such as those in stem cells niches. It will now be a challenge to further develop high-resolution live imaging of the CMA in these cells in order to test the proposed molecular control mechanisms.

In rapidly growing tissues, the sensitivity of katanin-mediated MT severing to tensile stress has been proposed as an additional array orienting mechanism [44,45]. This may explain the preferred orientation of cell division planes transverse to the main direction of growth in meristems, and potentially in later-stage embryos, where most divisions are also oriented orthogonal to the main growth axes. Explicit molecular hypotheses explaining katanin-mediated effects on array orientation have yet to be formulated, but the local modulation of severing activity can be readily implemented within our framework[32]. We envision that a combination of basic rules, which as we have argued here can only be unmasked in slow-growing cells, supplemented by moderating rules in rapidly growing cells will ultimately provide a complete picture of cell division control in plants, and means to modify these divisions for agricultural purposes.

## Acknowledgements

We thank Thomas Laux and Martin Hülskamp for critical reading of the manuscript. The work of BC was supported by a Wageningen University IPOP grant. The work of BMM is part of the research programme of the Netherlands Organisation for Scientific Research (NWO).

## Author Contributions

BC developed the computational framework and performed the simulations. BS and BM developed the questions and conceptual approach underlying the study. VW produced 3D stacks for WT and *clasp*, and analysed cell division phenotypes. TZ produced 3D stacks of *bdl* embryos and performed MT imaging in WT and *bdl*, and analysed the relation between MT, PPB and cell division orientation. CYL established the enabling technology for MT visualisation. BC, TZ, DW, BM and BS wrote the paper.

## Supplementary Materials

**Figure S1.**
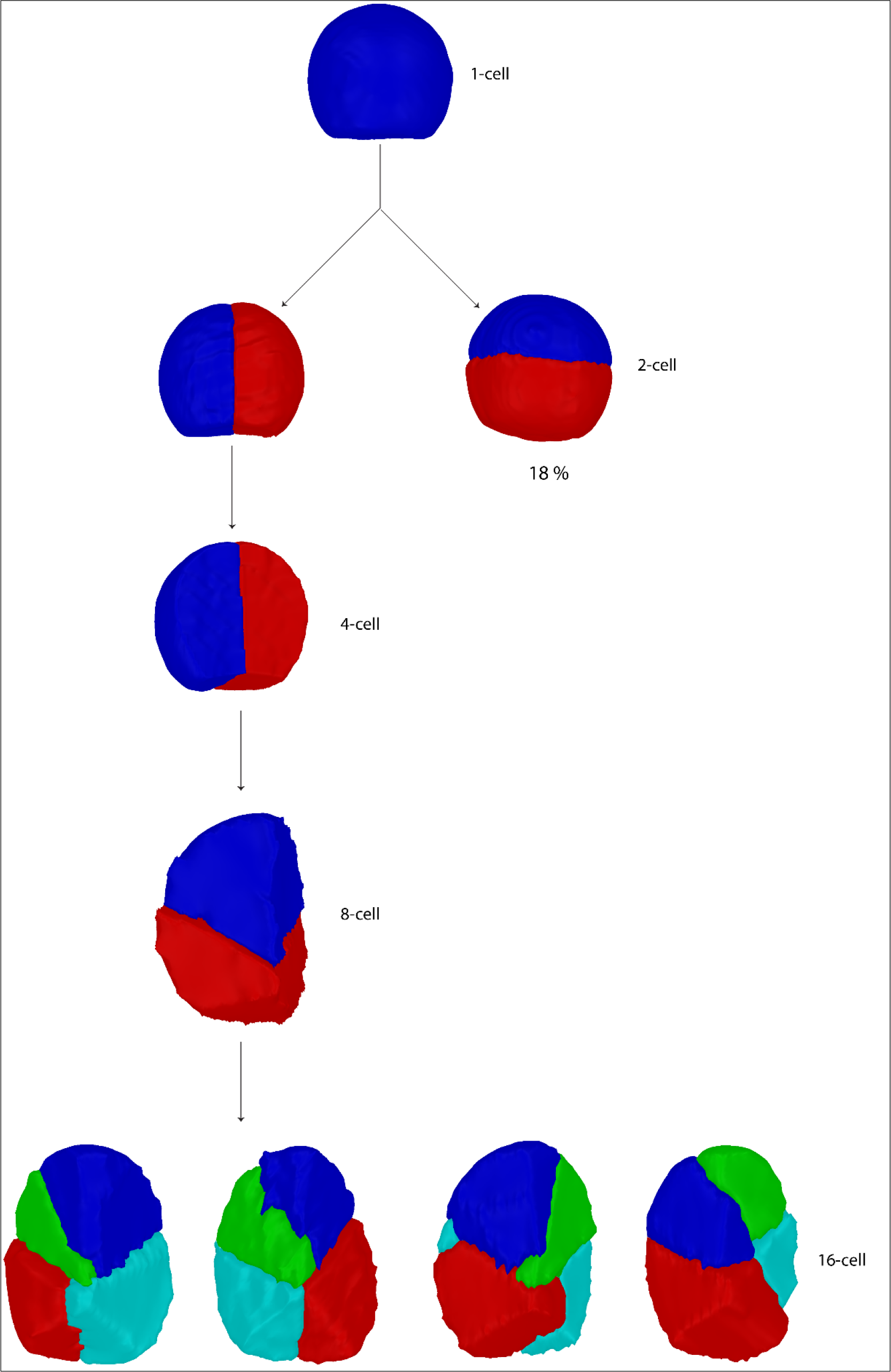
*Arabidopsis* early embryonic *bdl* division phenotype. A notable oblique division during 4- to 8-cell stage transition and aberrant divisions during 8- to 16 cell-stage transition. In analysis, 5 different embryos were used and only the correctly segmented cells were considered. We clearly observed these phenotypes, except the 18% horizontal divisions during 1-2-cell stage transition. However, such horizontal division phenotype has been reported in heterozygous *BDL/bdl* background[36].

**Figure S2.**
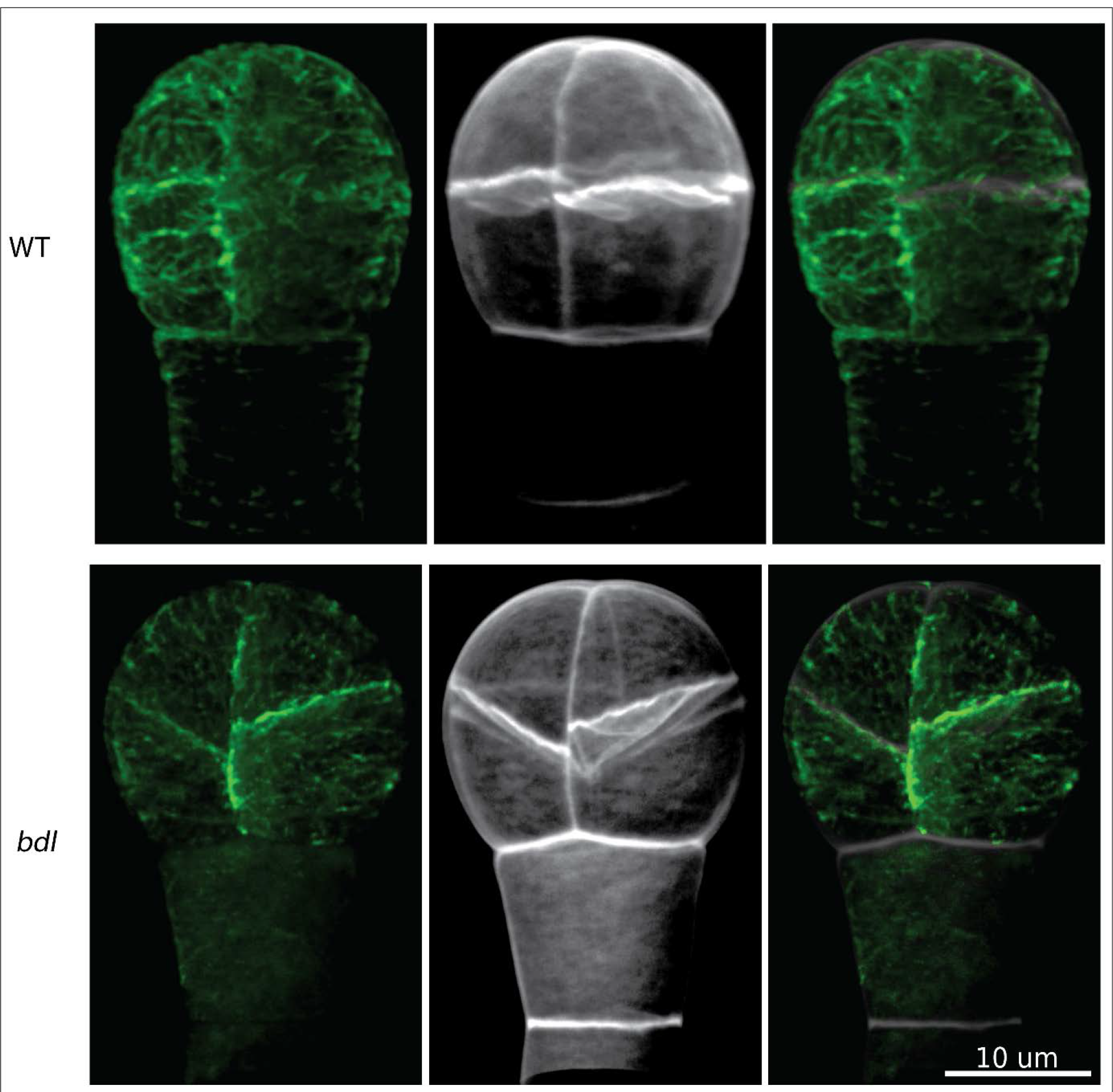
Cortical MT visualisation in *Arabidopsis* early embryonic cells using a pWOX2::TUA6-GFP reporter line in a RPS5A>>Col0 and RPS5A>>bdl background[47], revealing a shorter and sparser cortical MT network on *Arabidopsis* early embryonic cell faces in *bdl* mutant compared to WT.

**Figure S3.**
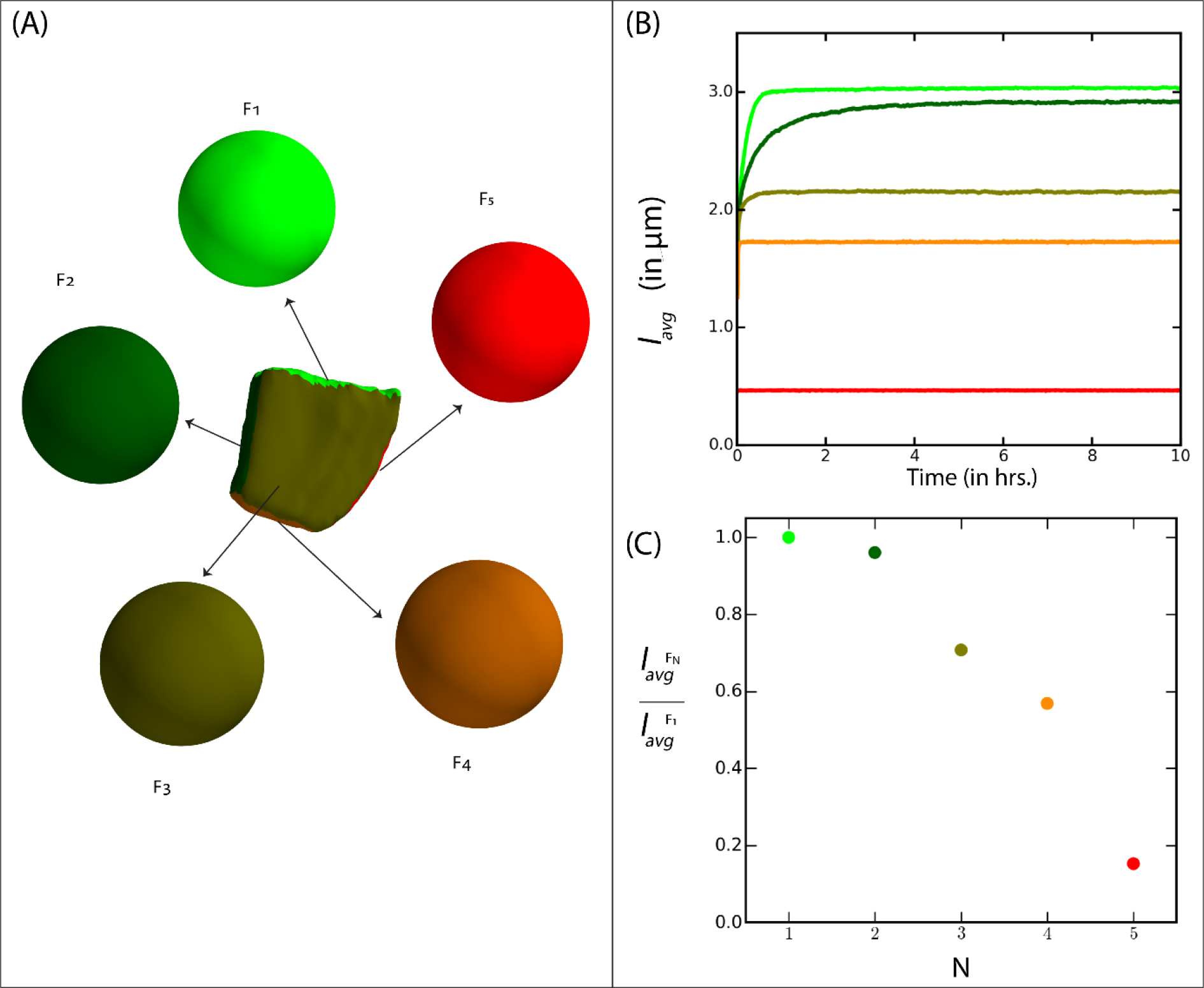
Simulation quantification of MT stability at developmentally different cell faces. (A) Schematic diagram of cell shape (a lower tier cell of 8-cell stage *Arabidopsis* embryo). This cell has five developmentally different faces (N =5), which we named as F_1_, F_2_, F_3_, F_4_ and F_5_ in the order of increased developmental age. We assigned the effect of increase in developmental age of a face (F) on the MT dynamics by an associated increase in the rate of dynamic catastrophe (r_c_^F^). An increase in rcF corresponds to a decrease in average MT length (*l_avg_*). Corresponding to each face, we created a sphere of radius ≈ 6 μm (average radius of an embryonic cell between 1- to 16-cell stage) with r_c_^F1^ < r_c_^F2^ < r_c_^F3^ < r_c_^F4^ < r_c_^F5^. (B) Simulation results of the steady state value of *l_avg_* corresponding to each of the spheres. (C) Fraction of steady state value of *l_avg_* at face F_N_ with respective to F_1_. This plot reflects the degree of MT stabilization at a developmentally old face (F_N_) compared to the newest face (F_1_).

**Figure S4.**
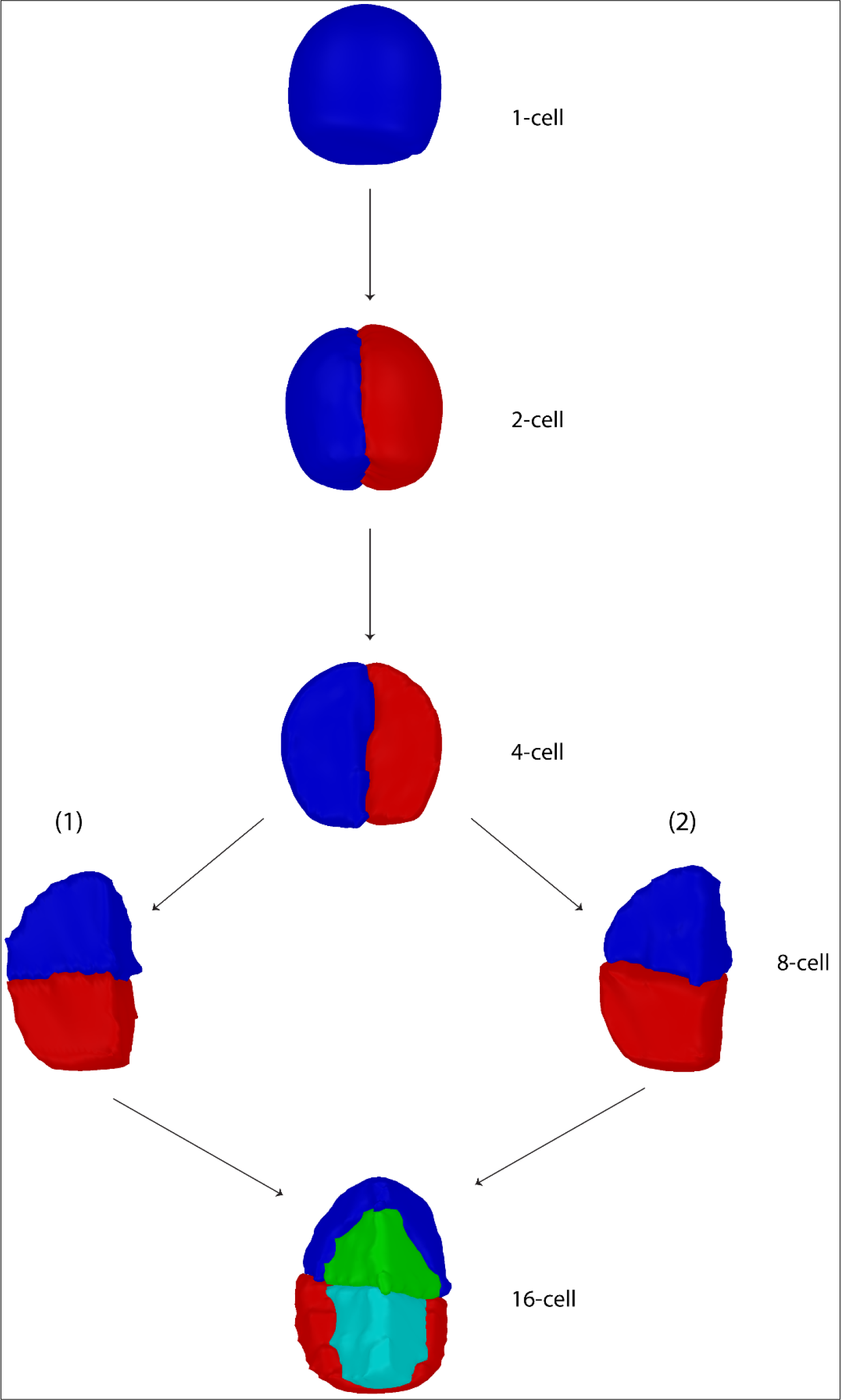
*Arabidopsis* early embryonic WT division phenotype. Notably, 4- to 8-cell stage transition reveals two types of division plane orientation phenotypes: (1) horizontal and (2) tilted horizontal division. Further quantification indicates that out of all observed divisions ≈ 46% (6/13) were horizontal and ≈ 54% (7/13) were tilted horizontal. Divisions during the 8- to 16- cell stage transition generate inner and outer tissue layers. In our analysis, 5 different embryos were used and only correctly segmented cells were considered.

**Figure S5.**
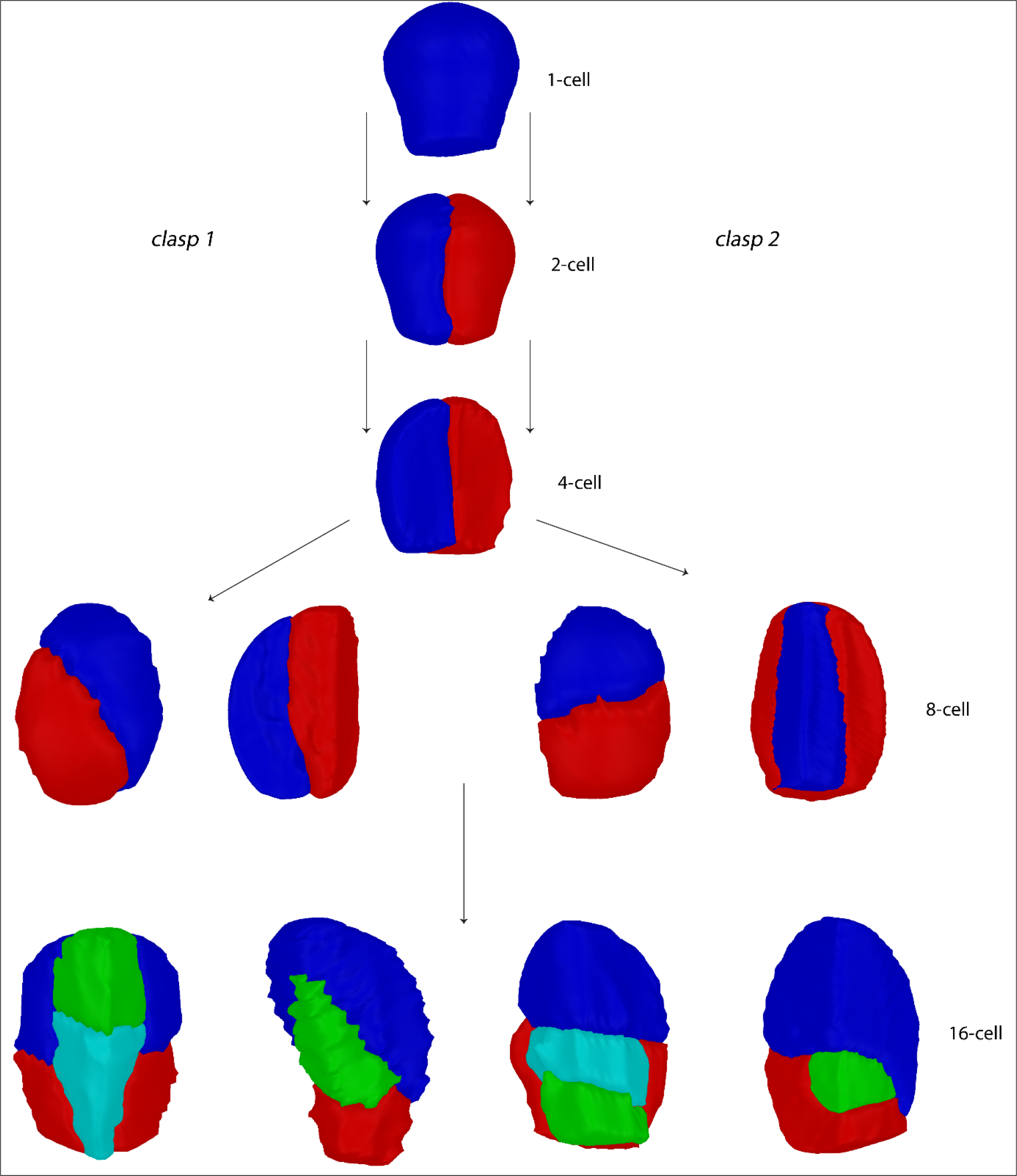
*Arabidopsis* early embryonic *clasp* division phenotype. Left branch corresponds to the *clasp 1* mutant allele and right to the *clasp 2* allele. Notably, 4- to 8-cell stage transition reveals division plane orientations which were skewed compared to WT. Out of all observed skewed divisions ≈ 65% (15/23) were oblique and ≈ 45% (8/23) generated the inward-outward layer. Further, 8- to 16-cell stage transition reveals aberrant division plane orientation.

**Figure S6.**
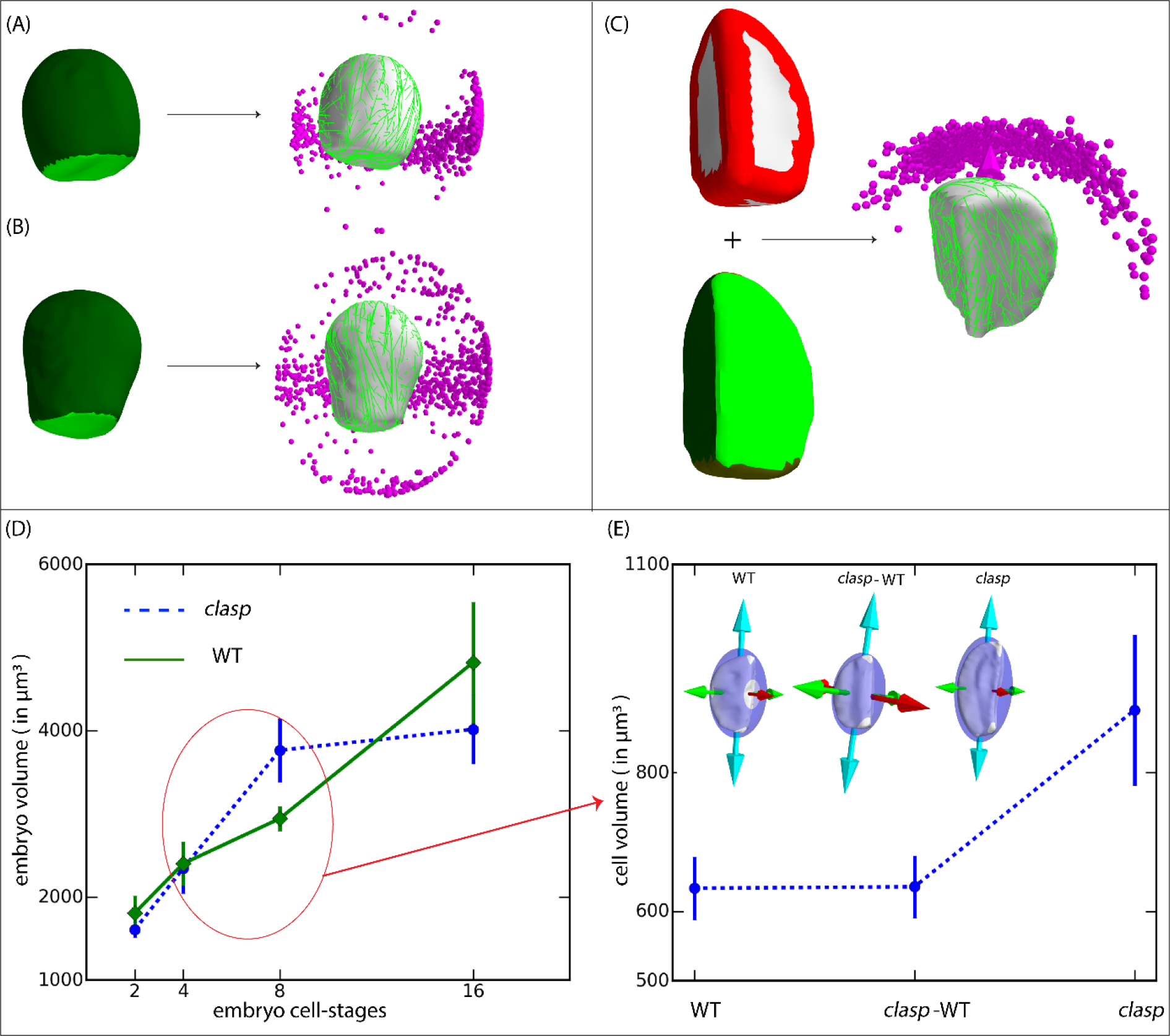
Simulating MT dynamics on 1-cell stage WT, 1- and 4-cell stage *clasp2* cell templates of *Arabidopsis* early embryo, and correlating the clasp division prediction with cell growth. For both (A) WT and (B) *clasp2*, MT dynamics was simulated by considering enhanced MT stabilization at the developmentally new basal cell face (green) than the apical face (dark green); see Figure S3 and Table S1. Predicted division was vertical which matched with the experimental observation but with full symmetry in the horizontal plane, i.e. a ring-shaped cluster of MT array orientation on the horizontal plane. The symmetry in MT array orientation actually reflects the default symmetry of these cell shapes. (C) MT simulation on an ‘original’ *clasp* 4- cell instead ‘reconstructed mother’ from 8-cell stage cell template. In these simulations, strong indication of a vertical division that creates inward-outward layer was revealed compare to rather weak prediction of such layering on merged cell templates (see Figure 5). (D) Comparison of embryonic volumes between WT (solid line) and *clasp (*mixture of *clasp 1* and *clasp 2* embryos, dashed line), during 2- to 16-cell stage transitions. (E) Comparative studies for changes in cell volume and cell shape anisotropy at the critical 4- to 8-cell stage transition in WT and *clasp* cells. At a given embryonic cell-stage not all cells were affected by loss of CLASP (i.e. *clasp* mutation), therefore we categorized the embryonic cells present in a *clasp* embryo into two sets: cells that showed normal WT like division phenotype were categorized as *clasp*-WT cells and the cells that gave typical *clasp* phenotype, i.e. formation of premature inner-outer layer at the 4- to 8-cell stage transition were categorized as (normal) *clasp* cells. Cell volume was calculated by re-creating about to divide 4-cell and therefore accounting post growth effect during 4- to 8-cell stage transition, by merging the corresponding 8-cell. Statistical analysis on cell volumes showed p < 0.05, indicating clear increase in cell volume in *clasp* affected cells than WT/*clasp*-WT cells. Further analysis via best fit ellipsoid showed a larger major to minor axis length ratio (k) in *clasp* (k ≈ 2.0) than WT/*clasp*-WT (k ≈ 1.7) cells, a possible reflection of growth induced change in shape anisotropy. This plot indicates in *clasp* already a significant growth in the average embryonic volume occurs during 4- to 8-cell stage transition, while in WT a significant growth in embryonic volume starts during 8- to 16-cell stage transition.

**Figure S7.**
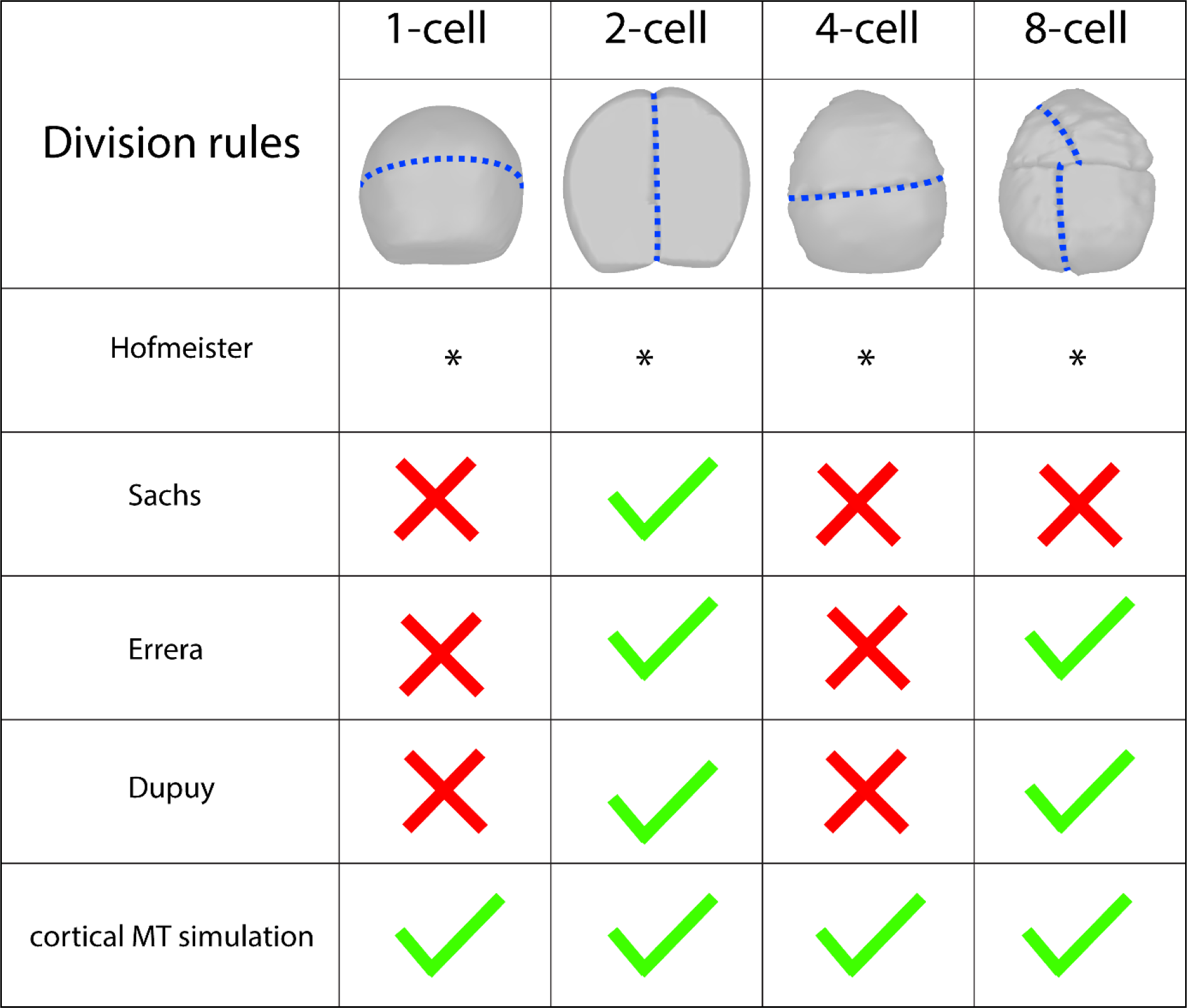
Comparison of the different rules of cell division when auxin signalling, a key genetic controller of *Arabidopsis* early embryonic cell division is compromised. Experimentally observed division plane orientations are shown by blue (dotted) lines. The symbols used in the table stands for: 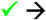 correct prediction (green), 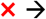 incorrect prediction (red) and * → not applicable, which holds uniformly for the Hofmeister rule, which is applicable for anisotropically growing cells only.

**Table S1.** Simulation parameterization in terms of average MT length (*l_avg_^F^*) at different cell faces (F) and edge-catastrophe multiplier (E_cat_) at the cell edges. For 1- to 8-cell stages transition of *Arabidopsis* early embryonic development, the number of available cell faces at a given embryonic stage is bounded in the range, 2 ≤ F ≤ 5, where 1-cell stage contains minimum number of faces (F = 2) and the lower tier of 8-cell stage contains the maximum number of faces (F = 5). Based on developmental age, we assigned a face identity (i.e F_1_, F_2_,F_3_,F_4_ and F_5_) and a degree of MT stabilization in terms of *l_avg_^F^* to each of these faces, where F_1_ is developmentally newest face and F_5_ is the oldest one (see Figure 4; Figure S3). Different combination of MT dynamics parameters were used[33] for calculating *l_avg_* to ensure formation of an ordered MT array on the default shape of cells taken from 1- to 8-cell stages. For any cell stage, 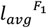 at F_1_ is calculated by using default shape simulation parameters and on other faces the MT dynamics catastrophe rate (r_c_) was tuned to vary 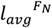 where N = 2,3,4,5. We tuned E_cat_ in the range, 0.0 ≤ E_cat_ ≤ 2.0. In all the figure panels: (1) for E_cat_ ≤ 0.5, the cell edges were coloured faint red indicating reduced edge-catastrophe, and (2) for E_cat_ > 0.5, the cell edges were coloured red indicating full edge-catastrophe or simply edge-catastrophe.

| Phenotype | Embryo stage transition | Average MT length (in $\mu\text{m}$) | Edge-catastrophe multiplier |
| --- | --- | --- | --- |
| WT | 1 → 2 cell | $l_{avg}^{F_2} \approx 0.9 \times l_{avg}^{F_1}$ | $E_{cat} \approx 0.5$ |
| | 2 → 4 cell | $l_{avg}^{F_2} \approx 0.9 \times l_{avg}^{F_1}$ and $l_{avg}^{F_3} \approx 0.7 \times l_{avg}^{F_1}$ | $E_{cat} \approx 0.5$ |
| | 4 → 8 cell | $l_{avg}^{F_2} \approx 0.9 \times l_{avg}^{F_1}$, $l_{avg}^{F_3} \approx 0.7 \times l_{avg}^{F_1}$ and $l_{avg}^{F_4} \approx 0.5 \times l_{avg}^{F_1}$ | $E_{cat} \leq 0.2$ |
| | 8 → 16 cell | $l_{avg}^{F_2} \approx 0.9 \times l_{avg}^{F_1}$, $l_{avg}^{F_3} \approx 0.7 \times l_{avg}^{F_1}$ and $l_{avg}^{F_4} \approx 0.1 \times l_{avg}^{F_1}$<br>$l_{avg}^{F_2} \approx 0.9 \times l_{avg}^{F_1}$, $l_{avg}^{F_3} \approx 0.7 \times l_{avg}^{F_1}$, $l_{avg}^{F_4} \approx 0.5 \times l_{avg}^{F_1}$ and $l_{avg}^{F_5} \approx 0.1 \times l_{avg}^{F_1}$ | $E_{cat} \leq 0.1$ |
| *bdl* | 1 → 2 cell | $l_{avg}^{F_2} \approx 0.9 \times l_{avg}^{F_1}$ | $E_{cat} \approx 0.5$ |
| | 2 → 4 cell | $l_{avg}^{F_1} = l_{avg}^{F_2} = l_{avg}^{F_3}$ | $E_{cat} \geq 0.5$ |
| | 4 → 8 cell | $l_{avg}^{F_1} = l_{avg}^{F_2} = l_{avg}^{F_3} = l_{avg}^{F_4}$ | $E_{cat} \geq 0.5$ |
| | 8 → 16 cell | $l_{avg}^{F_1} = l_{avg}^{F_2} = l_{avg}^{F_3} = l_{avg}^{F_4}$<br>$l_{avg}^{F_1} = l_{avg}^{F_2} = l_{avg}^{F_3} = l_{avg}^{F_4} = l_{avg}^{F_5}$ | $E_{cat} \geq 0.5$ |
| *clasp*<br>*clasp 1*<br>&<br>*clasp 2* | 1 → 2 cell | $l_{avg}^{F_2} \approx 0.9 \times l_{avg}^{F_1}$ | $E_{cat} \approx 0.5$ |
| | 2 → 4 cell | $l_{avg}^{F_2} \approx 0.9 \times l_{avg}^{F_1}$ and $l_{avg}^{F_3} \approx 0.7 \times l_{avg}^{F_1}$ | $E_{cat} \geq 1.0$ |
| | 4 → 8 cell | $l_{avg}^{F_2} \approx 0.9 \times l_{avg}^{F_1}$, $l_{avg}^{F_3} \approx 0.7 \times l_{avg}^{F_1}$ and $l_{avg}^{F_4} \approx 0.5 \times l_{avg}^{F_1}$ | $E_{cat} \geq 1.0$ |

## Materials and Methods

### Plant growth conditions

*Arabidopsis* seeds were surface-sterilized and dried seeds were subsequently sown on half-strength Murashige and Skoog (MS) medium. After a 48 hour cold treatment at 4 °C without light, seedlings were grown at 22 °C in standard long-day (16:8 h light:dark) growth conditions. After two weeks of growth, seedlings were transferred to soil and further grown under the same conditions. Siliques were harvested from mature plants for imaging. pRPS5A>>bdl embryos were generated by pollination of homozygous RPS5A-GAL4 pistils[48] with homozygous UAS-bdl[49] pollen. For *bdl* mutant MT-imaging, pWOX2::TUA6-GFP was crossed into the RPS5A-GAL4 line. Homozygous RPS5A-GAL4 X pWOX2::TUA6-GFP pistils were subsequently pollinated using either Col-0 or UAS-bdl.

### Staining and imaging conditions

**3D stacks**: *Arabidopsis* ovules of Col-0, *clasp 1* (SALK_120061) and *clasp 2* (SALK_83034) have been isolated from siliques in SCRI Renaissance 2200 solution on an objective slide[50,51]. The embryos have been popped out of the ovules by applying gentle pressure on the coverslip. 3D stacks of the embryos have been made using of Zeiss LSM710 with the 405nm laser. For generating 3D stacks of *bdl* embryos, we adapted the procedure described in Yoshida *et al.*[16].

**cortical MT**: *Arabidopsis* siliques were cut open using a fine needle and ovules were mounted and stained in a MT imaging solution (20% glucose, 10 uM Taxol, 0.1 M Pipes, pH 6.8, 1 mM EGTA, 1 mM MgSO4) containing 0.1% Renaissance (SR2200; Renaissance Chemicals; stock solution of the supplier was considered as 100%) cell wall counter-stain. Embryos were separated from ovules by gently pressing the coverslip. Images were taken within 30 minutes after release from the ovule. Sequential images were taken with 0.2 μm intervals to create high-resolution Z-stacks. Images were obtained using the Leica SP5 confocal laser scanning microscope equipped with photon-counting HyD detector and x60 water immersion objective. GFP and SR2200 signals were detected either at 488nm excitation/500-530nm emission or 405nm excitation/520-560nm emission wavelength, respectively.

### Cell segmentation and cortical MT projection

Cell segmentation and cortical MT projection are performed in MorphoGraphX (version 2.0)[46]. SR2200 stacks (tiff) were Guassian blurred using Sigma 0.3 μm. Blurred stacks were segmented using ITK watershed auto seeded segmentation with levels ranging from 500-1500. From segmented cells, meshes were created using Marching cubes 3D with Cube size in the range 0.5-1.0. cortical MT (pWOX2::TUA6-GFP) stacks (tiff) were loaded and overlaid with created corresponding cell meshes. Absolute cortical MT signal with a distance of 0-1 μm relative to the cell surface was projected on the cell mesh.

### Number of embryonic cells used in the simulations

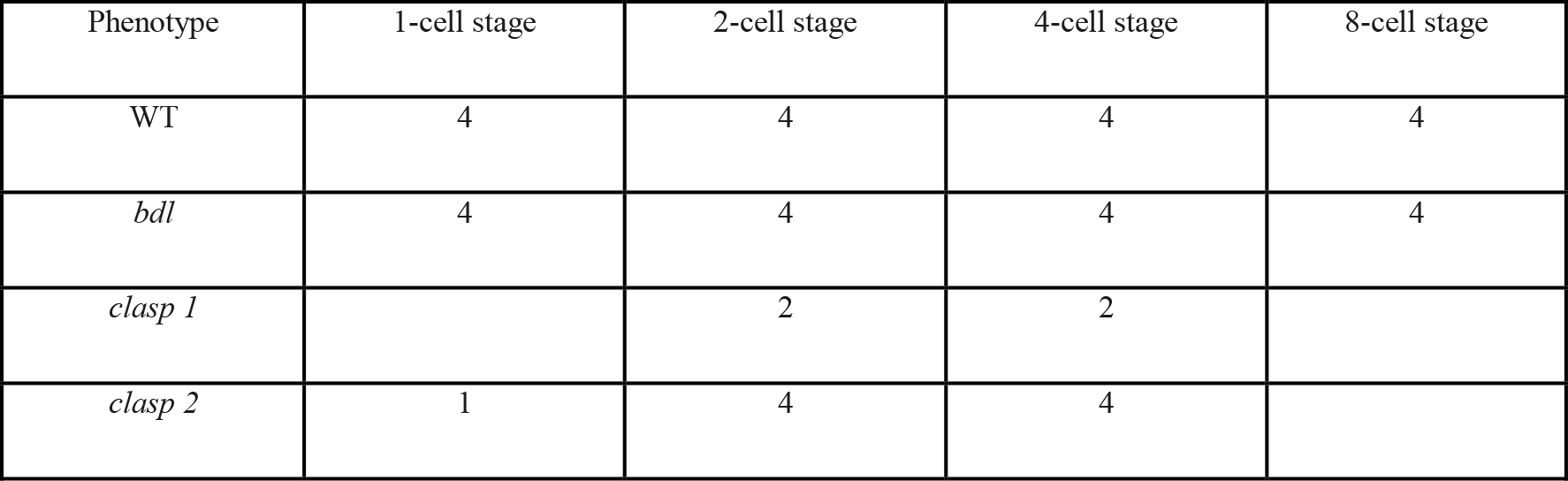

| Phenotype | 1-cell stage | 2-cell stage | 4-cell stage | 8-cell stage |
| --- | --- | --- | --- | --- |
| WT | 4 | 4 | 4 | 4 |
| *bdl* | 4 | 4 | 4 | 4 |
| *clasp 1* |  | 2 | 2 |  |
| *clasp 2* | 1 | 4 | 4 |  |

